# Phylogenetic analysis of the salinipostin γ-butyrolactone gene cluster uncovers new potential for bacterial signaling-molecule diversity

**DOI:** 10.1101/2020.10.16.342204

**Authors:** Kaitlin E. Creamer, Yuta Kudo, Bradley S. Moore, Paul R. Jensen

## Abstract

Bacteria communicate by small-molecule chemicals that facilitate intra- and inter-species interactions. These extracellular signaling molecules mediate diverse processes including virulence, bioluminescence, biofilm formation, motility, and specialized metabolism. The signaling molecules produced by members of the phylum Actinobacteria are generally comprised of γ-butyrolactones, γ-butenolides, and furans. The best known actinomycete γ-butyrolactone is A-factor, which triggers specialized metabolism and morphological differentiation in the genus *Streptomyces*. Salinipostins A-K are unique γ-butyrolactone molecules with rare phosphotriester moieties that were recently characterized from the marine actinomycete genus *Salinispora*. The production of these compounds has been linked to the 9-gene biosynthetic gene cluster *spt*. Critical to salinipostin assembly is the γ-butyrolactone synthase encoded by *spt9*. Here, we report the global distribution of *spt9* among sequenced bacterial genomes, revealing a surprising diversity of gene homologs across 12 bacterial phyla, the majority of which are not known to produce γ-butyrolactones. Further analyses uncovered a large group of *spt*-like gene clusters outside of the genus *Salinispora*, suggesting the production of new salinipostin-like diversity. These gene clusters show evidence of horizontal transfer between many bacterial taxa and location specific homologous recombination exchange among *Salinispora* strains. The results suggest that γ-butyrolactone production may be more widespread than previously recognized. The identification of new γ-butyrolactone biosynthetic gene clusters is the first step towards understanding the regulatory roles of the encoded small molecules in Actinobacteria.

**Importance:** Signaling molecules orchestrate a wide variety of bacterial behaviors. Among Actinobacteria, γ-butyrolactones mediate morphological changes and regulate specialized metabolism. Despite their importance, few γ-butyrolactones have been linked to their cognate biosynthetic gene clusters. A new series of γ-butyrolactones called the salinipostins was recently identified from the marine actinomycete genus *Salinispora* and linked to the *spt* biosynthetic gene cluster. Here we report the detection of *spt*-like gene clusters in diverse bacterial families not known for the production of this class of compounds. This finding expands the taxonomic range of bacteria that may employ this class of compounds and provides opportunities to discover new compounds associated with chemical communication.

## Introduction

Bacteria use chemical signaling molecules to regulate gene expression in a population dependent manner. This process, known as quorum sensing, controls group behaviors including swarming, bioluminescence, virulence, biofilm formation, cell competence, DNA uptake, public good production, and specialized metabolism. In many Gram-negative bacteria, quorum sensing is mediated by acyl-homoserine lactone (AHL) autoinducers (AIs) and their cognate receptors (1). In some Gram-positive bacteria, autoinducing peptides (AIPs) and their respective transmembrane two-component histidine sensor kinases control similar group behaviors (2). Among Actinobacteria, γ-butyrolactone signaling molecules regulate morphological development and specialized metabolite production. Given the importance of Actinobacteria for the production of antibiotics and other useful compounds, the discovery of new signaling molecules could facilitate the discovery of new natural products from the large number of “cryptic” gene clusters detected in actinomycete genome sequences.

To date, the types of signaling molecules known to be produced by Actinobacteria include γ-butyrolactones (3–25), γ-butenolides (26–30), furans (31, 32), PI factor (33), and N-methylphenylalanyl-dehydrobutyrine diketopiperazine (34) (Figure 1). Most of these were discovered from members of the genus *Streptomyces*. Sometimes referred to as actinobacterial “hormones”, signaling molecules are commonly produced in low amounts and have proven difficult to isolate and characterize. Many of these molecules not only induce the production of specialized metabolites, but also regulate bacterial morphogenesis and control complex regulatory systems (15). The first bacterial signaling molecule discovered was A-factor (autoregulatory factor, 2-isocapryloyl-3*R*-hydroxymethyl-γ-butyrolactone) from the actinomycete *Streptomyces griseus*. It was shown to trigger sporulation and the production of the antibiotic streptomycin (3). A-factor biosynthesis requires a γ-butyrolactone synthase and a reductase encoded by the genes *afsA* and *bprA,* respectively (15), and its elucidation revitalized the search to link signaling molecules to their biosynthetic genes (16, 32, 35–41). Most of the biosynthetically characterized γ-butyrolactones, γ-butenolides, and furans have been linked to *afsA* gene homologs via sequence similarity, biochemical verification, or A-factor receptor binding assays. However, many *afsA* gene homologs observed in *Streptomyces, Kitasatospora,* and *Amycolatopsis* genomes have yet to be linked to a small molecule (42–51). Likewise, the Acl series of γ-butyrolactones reported from *S. coelicolor* has not been linked to an AfsA homolog (3, 14, 18) (Figure 1).

**1.**
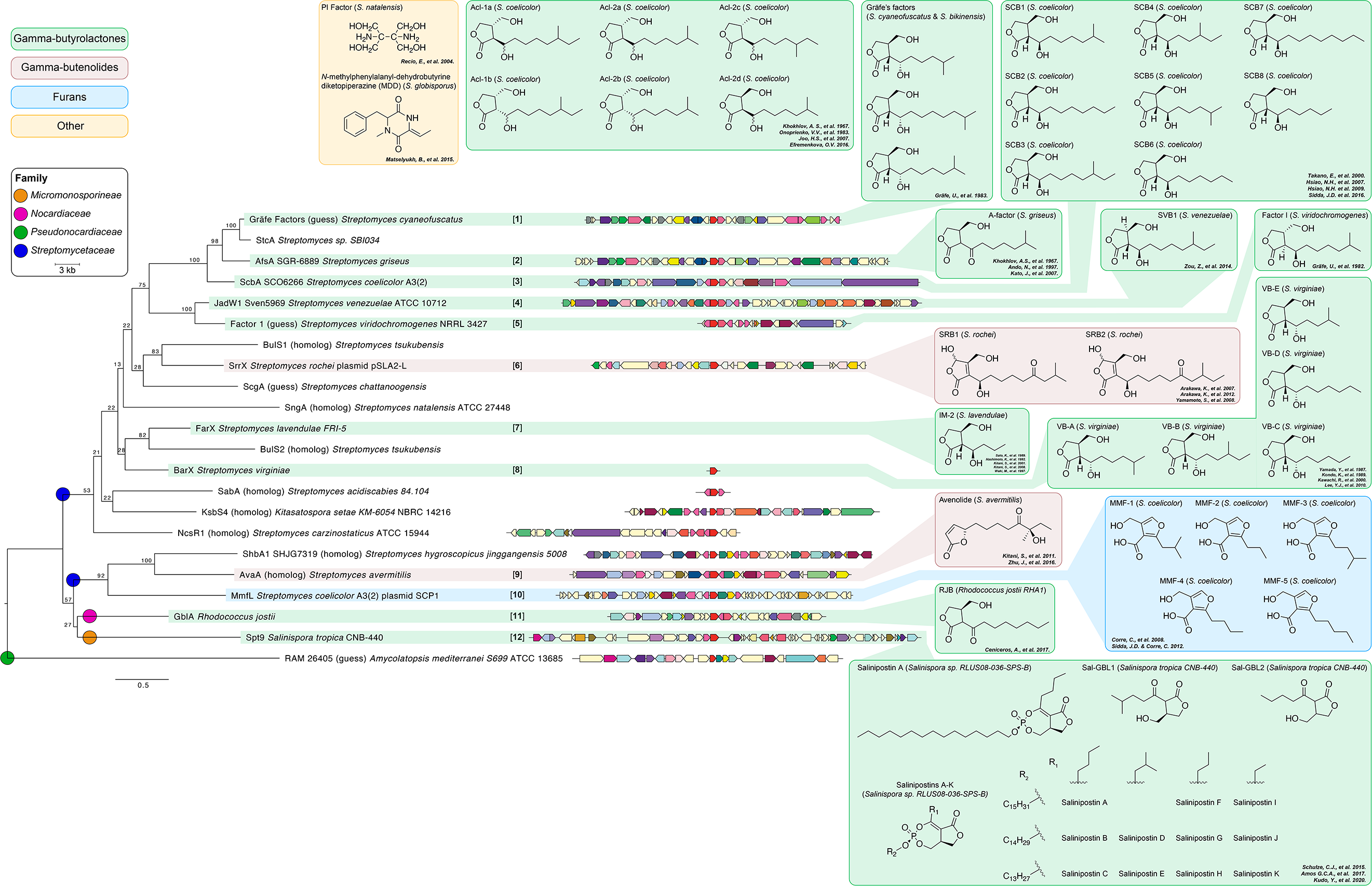
Actinobacterial AfsA homolog phylogeny, gene neighborhoods, and small molecule signaling products. Maximum likelihood phylogeny created with RaxML using a LG+I+G+F ProtTest model; branches are labeled with bootstrap support (500 replicates). Colored circles indicate Actinobacterial family. Gene neighborhoods are drawn 5’ to 3’ when genome sequences were available and aligned by AfsA homolog (red); other genes are colored by COG function. Colored boxes delineate γ-butyrolactones (green), γ-butenolides (red), and furans (blue). Those mapped to the tree have been experimentally linked to the gene cluster (references inset). Other actinobacterial signaling molecules not associated with AfsA are also shown (yellow). Bracketed numbers for each AfsA homolog and their associated signaling molecule product are referred to in subsequent figures.

While most A-factor-like molecules have been identified from *Streptomyces* strains, it remains possible that other actinobacterial taxa produce related signaling molecules. This includes the obligate marine actinomycete genus *Salinispora*, which is comprised of nine species: *Salinispora tropica, S. arenicola*, *S. oceanensis*, *S. mooreana*, *S. cortesiana*, *S. fenicalii*, *S. goodfellowii*, *S. vitiensis*, and *S. pacifica* (52, 53) isolated from marine sediments (54–57), seaweeds (55), and sponges (58, 59). The genus *Salinispora* has proven to be a prolific source of specialized metabolites (60) including the proteasome inhibitor salinosporamide A (61), which is currently in phase III clinical trials as an anticancer agent. Whole-genome sequencing of 118 *Salinispora* strains revealed 176 distinct biosynthetic gene clusters (BGCs), of which only 25 had been linked to their products (62). In a subsequent study, a majority of *Salinispora* BGCs were shown to be transcriptionally active under standard cultivation conditions, suggesting that many of their small molecule products were being missed using traditional detection and isolation techniques (63). Given that little is known about the regulation of specialized metabolism in this genus, it remains possible that quorum sensing signaling molecules may play a role in the regulation of BGC expression.

Recently, a series of compounds known as salinipostins A-K with rare bicyclic phosphotriesters were identified from a *Salinispora* sp. RLUS08-036-SPS-B (64). While these compounds were identified based on anti-malarial activity against *Plasmodium falciparum*, the γ-butyrolactone part of the salinipostin structure is reminiscent of *Streptomyces* A-factor (64, 65). Salinipostin biosynthesis was linked to the *spt* gene cluster via a knockout of the *afsA* homolog *spt9* in *S. tropica* CNB-440, which resulted in the elimination of salinipostin production (63). Subsequently, eight volatile bicyclic lactones, salinilactones A-H, were isolated and characterized from *S. arenicola* CNS-205 (66, 67). Two γ-butyrolactones Sal-GBL1 and Sal-GBL2 were also recently characterized from multiple *Salinispora* spp. strains (68). The Sal-GBLs, salinilactones A-H, and salinipostins A-K all share a bicyclic lactone motif and are proposed to originate from the same *spt* BGC (63, 66–68) (Supplementary Figure S1).

In this study, we set out to determine the distribution of *spt9* butyrolactone synthase homologs and the diversity of BGCs in which they reside, across all sequenced bacteria. We uncover that salinipostin-like BGCs are widely distributed outside of the genus *Salinispora* and exhibit gene rearrangements and some unusual gene fusions relevant to γ-butyrolactone biosynthesis. Finally, the evolutionary history of the salinipostin BGC indicates it was horizontally transferred between two *Salinispora* species at a location where they are known to co-occur.

## Materials and Methods

### Identification and distribution of Spt9 homologs

The predicted Pfam function of *spt9* was identified using the NCBI Conserved Domain Database prediction tool (69). AnnoTree (70) was used to determine the taxonomic distribution of the *spt9/afsA* Pfam03756 “A-factor biosynthesis hotdog domain-containing protein”. Protein-protein BLASTp (2.6.0+) (71) was used to query the *S. tropica* CNB-440 319-amino acid Spt9 protein sequence against the JGI IMG/MER sequence database (publicly available genomic sequence data integrated with JGI sequence data, all_img_core 2019) using an E-value cutoff of 1E-5 and at least 25% identity to identify the top 500 Spt9 homologs. To evaluate the gene neighborhood of the top matches, all but 33 *Salinispora* Spt9 homologs were removed, and a representative 20kb upstream and downstream of the Spt9 homologs were drawn; sequences were grouped into Actinobacterial family or Gammaproteobacterial class. Also included were 22 previously characterized *afsA* gene homologs including nine linked to the production of 34 γ-butyrolactone molecules; two linked to the production of three γ-butenolide molecules; and one linked to the production of five furan molecules.

To identify Spt9 homologs within the genus *Salinispora*, protein-protein BLASTp was used to search all publicly accessible *Salinispora* genomes with an E-value cutoff of 1E-5. PCR was used to confirm the integrity of the split *spt* BGC in *S. arenicola* CNS-296 and the presence of a hypothetical gene in *S. pacifica* CNS-143. PCR was performed by aliquoting 90 ng of gDNA into the PCR mixture consisting of 2X Phusion Green Hot Start II High-Fidelity PCR Master Mix (1.5 mM MgCl_2_, 200 μM each dNTP, 0.4 U Phusion enzyme; Thermo Scientific), 3% DMSO, and 0.5 μM of each forward and reverse primer (primer pair A: 6F [5’-ATCGAACGTGTCATCGAATGGC-3’], 6dntransR [5-CGTAGCCGAGGAAAGAAGCATC-3’]; primer pair B: 6F, 6dntrans-IGR_R [5’-TCGTTCATCAGAGGTCCCCTTC-3’]; primer pair C: 6F, 7R [5’-GATCAGATAGAGCATGGCGAGC-3’]. PCR conditions were as follows: primer pair A (6F, 6dntransR), 30 s of initial denaturation at 98°C, followed by 30 cycles of denaturation at 98°C for 5 s, annealing at 66°C for 20 s, and extension at 72°C for 30-50 s, followed by a final extension for 7 min at 72°C; primer pair B (6F, 6dntrans-IGR_R) same as previous but with annealing at 65.6°C for 20 s, and extension at 72°C for 69 s; primer pair C (6F, 7R), same as previous but with annealing at 65.7-66°C for 20 s, and extension at 72°C for 35-50 s, followed by a final extension for 5-7 min at 72°C. The resulting products were visualized on a 0.8% agarose gel run in 1X TAE at 95-97V for 30-60 min; excised, purified, and Sanger sequenced in forward and reverse directions (Eton Bioscience, San Diego, CA), trimmed, and mapped to their respective genomes in Geneious v8.1.9 (72).

### Identification of salinipostin-like BGCs

ClusterScout (73) searches were performed to identify salinipostin-like BGCs in other sequenced genomes using the following Pfam functions: *spt1* Pfam00391, Pfam01326; *spt2* Pfam00501; *spt4* Pfam00550; *spt5* Pfam07993; *spt6* Pfam00334; *spt7* Pfam01040; *spt8* Pfam00296; *spt9* Pfam03756. It should be noted that antiSMASH v4 and v5 (74, 75) do not fully identify *spt1-2* in the salinipostin butyrolactone BGC, thus other methods were used to find *spt-*like BGCs. Independent searches were run with a minimum requirement of either 3, 4, or 5 Pfam matches, a maximum distance of <10,000bp between each Pfam match, and a minimum distance of 1bp from the scaffold edge. The boundaries of each match were extended by a maximum of 10,000bp to help identify full biosynthetic operons. For some searches, the *spt9*/*afsA* Pfam was defined as essential. MultiGeneBlast (76) was also used to query the contiguous *S. tropica* CNB-440 salinipostin *spt1-9* gene cluster against the NCBI GenBank Bacteria BCT subdivision database. Finally, the STRING v11 database (77) was queried using Spt1-9 to identify significant protein-protein interactions, gene neighborhoods, and gene co-occurrences within 5,090 organisms. Biosynthetic clusters retrieved from each ClusterScout, MultiGeneBlast, and STRING search were manually inspected for *spt* Pfams and gene organization.

### Phylogenetic distribution of Spt9 homologs and salinipostin-like BGCs

A maximum likelihood amino acid phylogeny was generated from the top 403 Spt9 homologs and an additional 22 experimentally characterized Spt9/AfsA proteins. The sequences were aligned with MUSCLE (78) within the Mesquite system for phylogenetic computing (79) and analyzed using ProtTest 3.4.2 (80) to determine an amino acid model for tree calculations. RAxML (81) was used to create a tree using ML + rapid bootstraps with 500 replicates. A second phylogeny was generated for the Spt9 homologs observed in salinipostin-like BGCs using the same parameters. The topologies of these trees and branch support were confirmed using PhyML (82) with SMS Smart Model Selection (AIC model selection; BIONJ tree searching, NNI tree improvement, and an aLRT SH-like fast likelihood method) (83).

To test if the *Salinispora* salinipostin BGC was acquired as an intact gene cluster, Spt1-9 proteins from 116 *Salinispora* genomes were aligned with MUSCLE (78) and PhyML (82) used to calculate a phylogenetic tree for each protein with automatic SMS Smart Model Selection (AIC model selection; BIONJ tree searching, NNI tree improvement, and an aLRT SH-like fast likelihood-based method) (83). The nine Spt1-9 protein trees were compared for congruency with a concatenated *Salinispora* species tree created using the following 11 single-copy protein sequences: DnaA, GyrB1, GyrB2, PyrH, RecA, Pgi, TrpB, AtpD, SucC, RpoB, and TopA as previously reported (84). Spt1-9 protein sequences were also concatenated (3,758 total amino acid characters), aligned with MUSCLE (78), and a maximum likelihood tree calculated using SMS and PhyML with the previously described parameters.

FigTree (85) and the Interactive Tree of Life (iTOL v4) (86) were used to visualize phylogenetic trees and Actinobacterial families were assigned using a recently proposed phylogeny (87). Fused genes consisting of functional domains from two Spt proteins were identified using Geneious v8.1.9 (72).

## Results

### Taxonomic distribution of Spt9 homologs

The γ-butyrolactone synthase AfsA is critical in the biosynthesis of the *Streptomyces* signaling molecule A-factor (15). The identification of the *afsA* homolog *spt9* in the salinipostin (*spt*) BGC and its role in catalyzing the γ-butyrolactone ring formation in salinipostin biosynthesis led us to explore the distribution of *spt9/afsA* homologs among sequenced bacterial genomes. We first confirmed that Spt9 contains two AfsA-like hot-dog fold superfamily domains that are distantly related to the FabA and FabZ β-hydroxyacyl-ACP dehydratases associated with fatty-acid biosynthesis in *Escherichia coli* (88). Using AnnoTree (70), 1,230 *spt9/afsA* Pfam03756 hits were identified out of the 27,000 reference genomes in the Genome Taxonomy Database (Figure 2). Surprisingly, these were distributed among 12 phyla, the majority of which are not known for the production of γ-butyrolactone signaling molecules. The phylum Actinobacteria contained 74% of the hits, Proteobacteria had 21% of the hits, and the rest were scattered across 10 additional phyla. Noticeably, of the 3,579 Actinobacteria in the reference Genome Taxonomy Database, 25% (911) contained the AfsA Pfam03756 compared to only 3% (256) of the 8,882 Proteobacteria.

**2.**
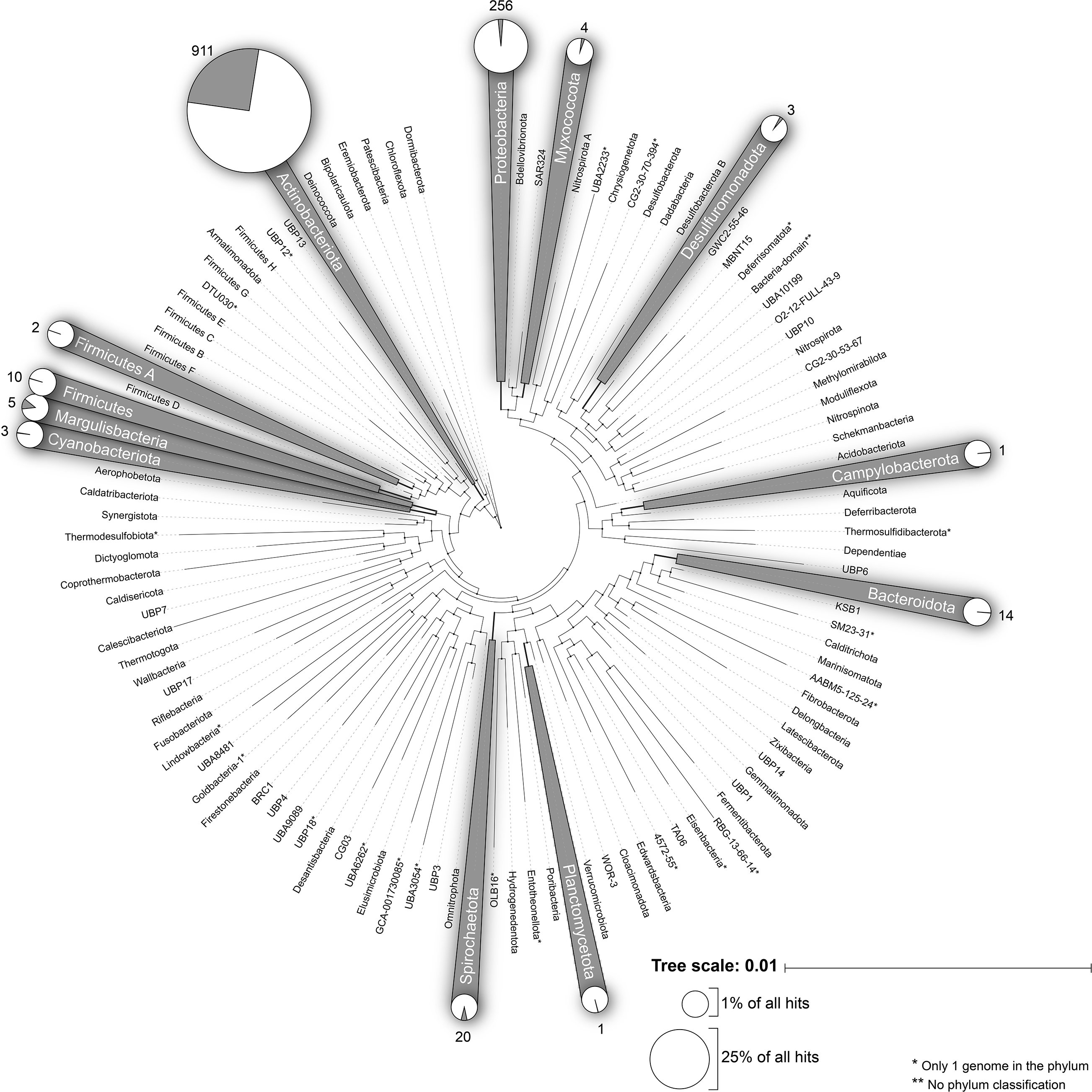
Distribution of Spt9 homologs across 27,000 bacterial genomes. Shaded taxa contain Spt9/AfsA Pfam03756 A-factor biosynthesis hotdog domain-containing protein homologs as determined using AnnoTree. Pie charts show the proportion of genomes in that taxon with a Spt9 homolog, with the total number of hits indicated. Pie chart size is proportional to the percentage of Spt9/AfsA hits out of the 1,230 detected across all taxa.

### Spt9 phylogeny and gene environment

To assess the detailed phylogenetic relationships of the Spt9 homologs, we conducted a BLASTp search to identify the top related Spt9 proteins across the ~70,000+ bacterial genomes in the 2019 JGI IMG Blast database (89). The top 500 homologs shared at least 25% amino acid identity with Spt9, of which the top 403 were further analyzed after removing duplicate *Salinispora* Spt9 homologs. We additionally identified 22 AfsA homologs that have been experimentally linked to the production of a diverse array of γ-butyrolactones, γ-butenolides, and furans (Figure 1). A maximum likelihood phylogeny generated using these Spt9/AfsA homologs was incongruent with the established taxonomic relationships of the strains in which the sequences were detected (Figure 3, Supplementary Figure S2). One prominent example includes the *Salinispora* Spt9 sequences, which are sister to a homolog in *Streptomyces phaeofaciens*. These sequences fall within a larger clade comprised of diverse members of the Gammaproteobacteria (*Citrobacter koseri* and *Dickeya sp.*) and Actinobacteria (*Cellulomonas cellasea* and *Rhodococcus* sp.) as opposed to cladding with other members within the *Micromonosporaceae* family. The 22 AfsA-homologs that have been linked to the production of γ-butyrolactones, γ-butenolides, and furans are restricted to one large clade in the phylogeny and distinct from the northern end of the tree, which contains the *Salinispora* Spt9 sequences (Figure 3).

**3.**
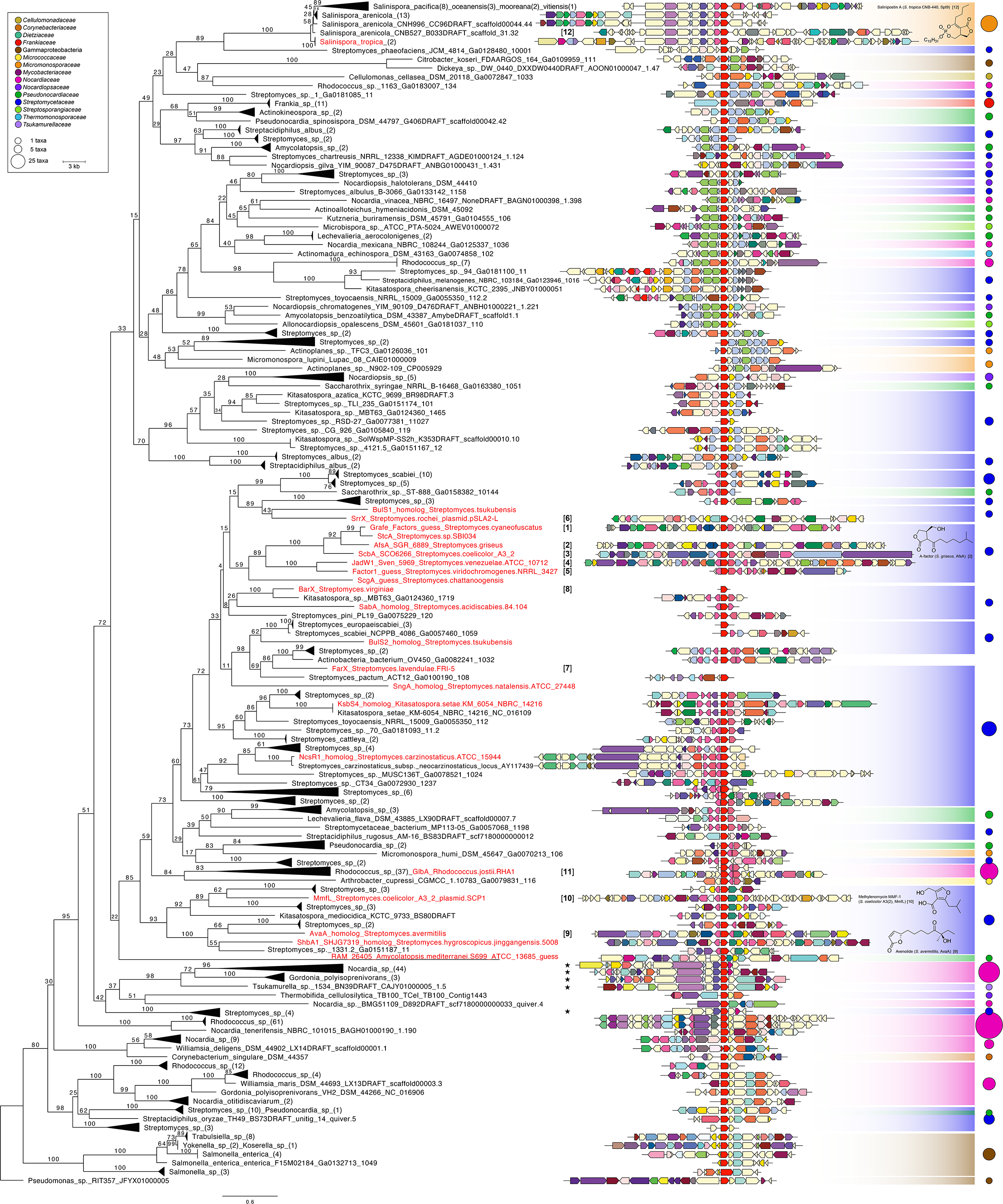
Phylogeny and gene environments of characterized AfsA and top Spt9 homologs. Condensed maximum likelihood phylogeny of the top Spt9 homologs (403, black) and 22 experimentally characterized AfsA homologs (22, red) linked to known molecules. The tree was calculated with a WAG+I+G+F ProtTest model with 500 replicates in RaxML; branches are labeled with bootstrap support. Taxonomically coherent clades are collapsed with the number of sequences indicated in parentheses. The *Pseudomonas sp.* RIT357 AfsA homolog was used as an outgroup. Gene neighborhoods are drawn 5’ to 3’ and aligned with the Spt9 homolog (red); genes are colored by COG function as annotated by JGI IMG/MER. Shaded rectangles indicate Actinobacterial family or Gammaproteobacterial class (see legend) with circles proportional to the number of sequences in each familial clade. Representative chemical structures are shown (γ-butyrolactones: salinipostin A from *Salinispora tropica* CNB-440, A-factor from *Streptomyces griseus*; γ-butenolide: avenolide from *Streptomyces avermitilis*; furan: methylenomycin from *Streptomyces coelicolor* A3(2)) and bracketed numbers correspond to the AfsA homologs and their associated signaling molecule products in Figure 1. Stars indicate salinipostin-like BGCs.

The large number of Spt9/AfsA homologs illustrates considerable potential for the discovery of new γ-butyrolactone synthase mediated chemical diversity (Figure 3). New biosynthetic routes are supported by the diverse gene environments in which these Spt9/AfsA homologs are observed. For example, antiSMASH 5 (75) analyses reveal that some Spt9 homologs are close to genes encoding ketosynthase and thiolase enzymes, suggesting they are part of larger PKS gene clusters. Among *Salinispora* strains, six Spt9 homologs were observed outside of the *spt* BGC. Two of these were observed in *Salinispora oceanensis* strains CNT-124 and CNT-584, each of which contain two Spt9 homologs in a type II PKS BGC in addition to the *spt* BGC. The other four were observed in *Salinispora fenicalii* strains CNT-569 and CNR-942, which lack the *spt* BGC but instead have a Spt9 homolog both in a type II PKS BGC and a butyrolactone non-ribosomal peptide synthetase (NRPS) BGC.

Despite incongruence with the species phylogeny, the gene environments surrounding some Spt9/AfsA homologs show evidence of gene conservation. For example, in the clades near the characterized Spt9/AfsA homologs, many *Streptomyces* and *Kitasatospora* species showed a general conservation of a *bprA* gene homolog (dark pink) next to *spt9* homologs as required for A-factor biosynthesis in *Streptomyces griseus* (Figure 3). The three Spt9 homologs that are closest to the *Salinispora* Spt9 sequence (observed in *Streptomyces phaeofaciens, Dickeya sp.* and *Citrobacter koseri*) contain a *spt4* acyl carrier protein homolog (pale blue). Below the *Salinispora* Spt9 clade, conservation of two 3-oxoacyl (acyl-carrier-protein) synthases (light green), an acyl carrier protein (light blue*, spt4* homologs), a hydrolase (pale yellow), and a 3-oxoacyl-(acyl-carrier-protein) reductase (light blue) is observed across taxonomically diverse *Streptomycetaceae*, *Nocardiopsaceae*, *Nocardiaceae*, *Thermomonosporaceae*, *Pseudonocardiaceae*, and *Streptosporangiaceae* strains. At the bottom of the tree, the Gammaproteobacterial outgroup *Pseudomonas sp.* RIT357 shares conserved genes around the Spt9 homolog with other diverse families of bacteria including other Gammaproteobacteria, *Nocardiaceae*, *Streptomycetaceae*, and *Pseudonocardiaceae* with a putative hydrolase of the HAD superfamily (light yellow), a cytochrome P450 (light teal), and a MFS (major facilitator superfamily) protein transporter (light yellow). The presence of a few conserved genes near Spt9 homologs yet the lack of fully syntenic Spt9 gene neighborhoods suggests there are some functional similarities involving Spt9 homologs across diverse bacterial families. The observation of diverse gene environments flanking Spt9 homologs illustrate the enormous potential for γ-butyrolactone, γ-butenolide, and furan production among bacteria not known to produce these molecules through potentially new biosynthetic routes.

### Targeted search for spt-like BGCs

A number of Spt9 gene neighborhoods outside of *Salinispora* caught our attention due to similarities with the salinipostin BGC (Figure 3). To search more thoroughly for *spt*-like BGCs among sequenced genomes, we searched for *spt-*like BGCs using ClusterScout (73), MultiGeneBlast (76) and STRING v11 (77). These efforts led to the identification of 91 *spt*-like BGCs spanning six actinomycete families within the genera *Nocardia*, *Gordonia*, *Tsukamurella*, *Mycobacterium*, *Dietzia, Streptomyces*, *Kitasatospora, Rhodococcus,* and *Kutzneria* (Figure 4). All of these BGCs possess *spt1, spt5, spt6, spt7,* and *spt9* homologs with *spt9* towards the 3’ end of the cluster as seen in *Salinispora*. Notably, none of the *spt*-like BGCs contain the flavin-dependent oxidoreductase *spt8*, whose role is unknown in salinipostin biosynthesis. None of these *spt*-like BGCs have been linked to the small molecules they encode.

**4.**
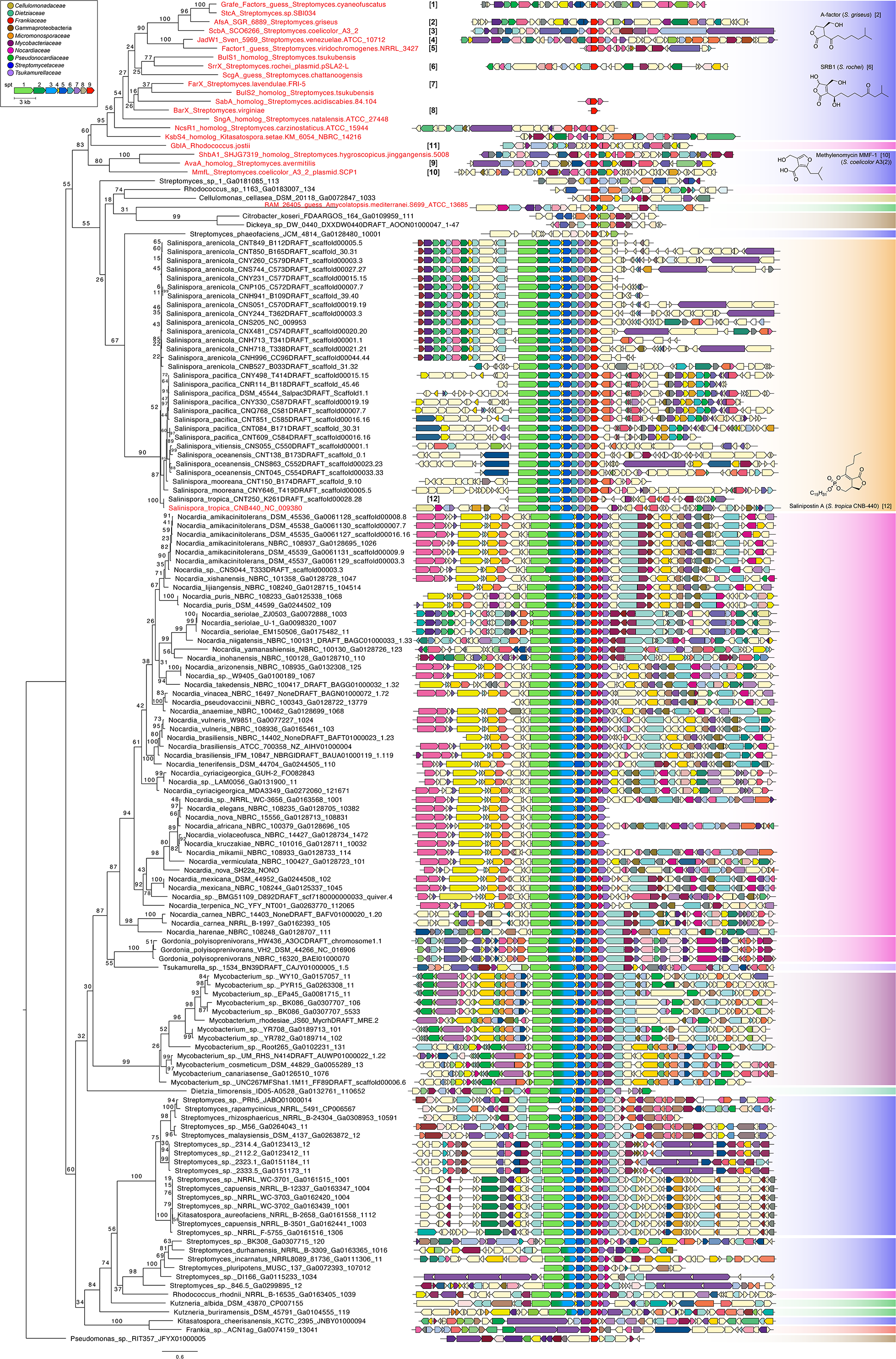
Phylogeny of Spt9 homologs within salinipostin-like BGCs. The maximumlikelihood phylogeny was calculated with a WAG+I+G+F ProtTest model with 500 replicates in RaxML; branches are labeled with bootstrap support. Gene neighborhoods are drawn 5’ to 3’ and aligned with the Spt9 homolog (red). The names of Spt9/AfsA homologs linked to the production of specific compounds are colored in red. Gene fusions are shown with approximate transition points and neighboring genes are colored by COG function as annotated by JGI IMG/MER. Colored rectangles indicate Actinobacterial family or Gammaproteobacterial class (see legend). Representative chemical structures are shown (γ-butyrolactones: A-factor from *Streptomyces griseus,* salinipostin A from *Salinispora tropica* CNB-440; γ-butenolide: SRB-1 from *Streptomyces rochei*; furan: methylenomycin MMF-1 from *Streptomyces coelicolor* A3(2)) and bracketed numbers correspond to the AfsA homologs and associated signaling molecule products in Figure 1.

A maximum likelihood phylogeny of the Spt9 homologs observed in *spt*-like BGCs clearly delineates them from the 11 AfsA homologs linked to γ-butyrolactone, γ-butenolide, and furan biosynthesis (Figure 4). Compared with the nine gene salinipostin BGC identified in *Salinispora* (Supplementary Figure S1), we observed gene reorganizations and fusions in other bacteria (Figure 4). Most notably, *spt2* and *spt3* are fused across the large clade bracketed by *Nocardia* and *Dietzia timorensis*, as well as two *Kutzneria* species and five *Streptomyces* species at the most southern part of the tree. This fusion is conserved across most BGCs except those observed in *Salinispora*, four *Streptomyces* species, and *Rhodococcus rhodnii* NRRL B-16535. A second example of gene fusion is observed between *spt6* and *spt9* in *Dietzia timorensis*. Alignment of the *spt2, spt3, spt6,* and *spt9* fused and individual genes reveals maintenance of the functional domains (Supplementary Figure S3). These gene fusions, also known as Rosetta gene fusions (90, 91), suggest a functional interaction between the encoded proteins in the biosynthesis of salinipostin-like γ-butyrolactone molecules. The gene fusions could have arisin from the single-domain individual *spt2, spt3, spt6,* and *spt9* genes in the *Salinispora spt* BGC and thus suggest some evolutionary selective advantage for these specific co-localized biosynthetic genes to become fused.

Many of the *spt*-like BGCs differed in gene order compared to that observed in *Salinispora* while others contained extra genes in the cluster including a nitroreductase (dark pink). It is noteworthy that *spt8*, a flavin-dependent oxidoreductase (Supplementary Figure S1) was unique to the *Salinispora spt* BGC; however, no definitive function of *spt8* has been suggested in the proposed biosynthesis of the salinipostins (68) or salinilactones (66, 67). Similarly, some *Streptomyces spt*-like BGCs did not contain *spt2* (AMP-ligase) and *spt4* (acyl carrier protein) homologs, which are proposed to help load and carry the R_2_ aliphatic sidechain during salinipostin biosynthesis, respectively (68). These observations suggest additional structural diversity remains to be discovered. Furthermore, two BGCs found in *Kitasatospora cheerisanensis* and *Frankia sp.* contained *spt9* and *spt2* homologs next to a type I polyketide synthase (PKS), suggesting a potential role in the regulation of the PKS BGC as observed in methylenomycin biosynthesis (31). Overall, we observed that some of the 91 *spt*-like BGCs occur in different genomic locations within the same genus, which could support BGC migration or horizontal gene transfer. However, there is also evidence of vertical inheritance based on gene conservation in some strains. To investigate this further, we focused on the genus *Salinispora*, where BGC migration and transfer events have been previously reported (62).

### The spt BGC in the genus Salinispora

The salinipostin BGC (*spt1-9*) is highly conserved within the genus *Salinispora* (62). Notably, only five strains have been identified via genome mining that lack the *spt* BGC and these belong to the recently described species *S. fenicalii*, *S. goodfellowii*, and *S. vitiensis* (Supplementary Figure S4A). At the species level, the *spt* BGC is commonly observed in the same genomic environment (Supplementary Figure S5) within previously defined genomic islands (62, 92). For example, the *spt* BGC is located in Genomic Island (GI) 20 in *S. tropica* (62) with notable conservation of upstream and downstream regions. Similarly, positional conservation of the *spt* BGC in Genomic Island 15 (62) is observed in most *S. arenicola* and *S. pacifica* strains.

Variations in the *spt* BGC are observed in three *Salinispora* strains (Figure S5). In the case of *S. arenicola* CNS-296, *spt1-6* and *spt7-9* are split onto different contigs and flanked by transposases. Targeted PCR amplifications of *spt6* and the neighboring transposase confirmed that the BGC is indeed split (Supplementary Figure S6A). Attempts to amplify a region between *spt7* and the downstream hypothetical gene resulted in multiple PCR products likely due to multiple copies of the hypothetical gene in the *S. arenicola* CNS-296 genome and was thus uninformative. However, primers spanning *spt6-7* resulted in a product that was 3.2kb larger than what was observed from a contiguous BGC in *S. arenicola* CNQ-884 (Supplementary Figure S6B). Sequences obtained from the ends of this amplicon mapped poorly to *spt6* (67% identity) and better to *spt7* (99% identity), providing additional support for an insertion between *spt6-7* in the *S. arenicola* CNS-296 *spt* BGC. It remains to be determined if *S. arenicola* CNS-296 produces salinipostins. In contrast, the detection of *spt* genes on multiple contigs in *S. arenicola* CNT-088 and *S. pacifica* CNS-143 is likely due to poor genome assembly (93). We also investigated the hypothetical gene annotated between *spt6-7* in *S. pacifica* CNS-143. A comparison of PCR products spanning *spt6-7* in this strain with *S. arenicola* CNQ-884, where the *spt* BGC is contiguous, yielded amplicons of the same size and with the same conserved *spt6-7* domains (Supplementary Figure S6B). This indicates that the extra hypothetical gene called in *S. pacifica* CNS-143 is likely an error and provides further support for conservation of the *spt* BGC in the genus *Salinispora*.

While conservation of the *spt* BGC within *Salinispora* supports vertical inheritance, an Spt9 phylogeny (Supplementary Figures S4B, S5) reveals incongruencies with the established *Salinispora* species phylogeny (52, 62) (Supplementary Figure S4A)that are consistent with recombination events. In one example, Spt9 sequences from *S. vitiensis* CNS-055 and *S. cortesiana* CNY-202 occur within the *S. oceanensis* clade as opposed to outside of it (Supplementary Figures S4, S5). A more pronounced example is the placement of *S. tropica* Spt9 sequences within the larger *S. arenicola* clade, suggesting the former acquired its sequence from the latter. This recombination event appears to have involved the entire BGC since the remaining *S. tropica* Spt1-8 sequences share a similar phylogeny (Figure S4B). As predicted, a concatenated phylogeny of Spt1-9 (Figure 5A) is incongruent with the *Salinispora* species phylogeny (Supplementary Figure S4A) and supports a horizontal exchange of the BGC between *S. arenicola* and *S. tropica*. Mapping the geographic origin of the strains onto the tree reveals that all twelve *S. tropica* and the six most closely related *S. arenicola* sequences all originated from ocean sediments collected in the Bahamas and the Yucatán (Figure 5B). A closer examination reveals that the two species co-occur at four of the five collection sites (Figure S7). This geographic proximity would provide opportunities for BGC horizontal gene transfer and homologous recombination to occur.

**5.**
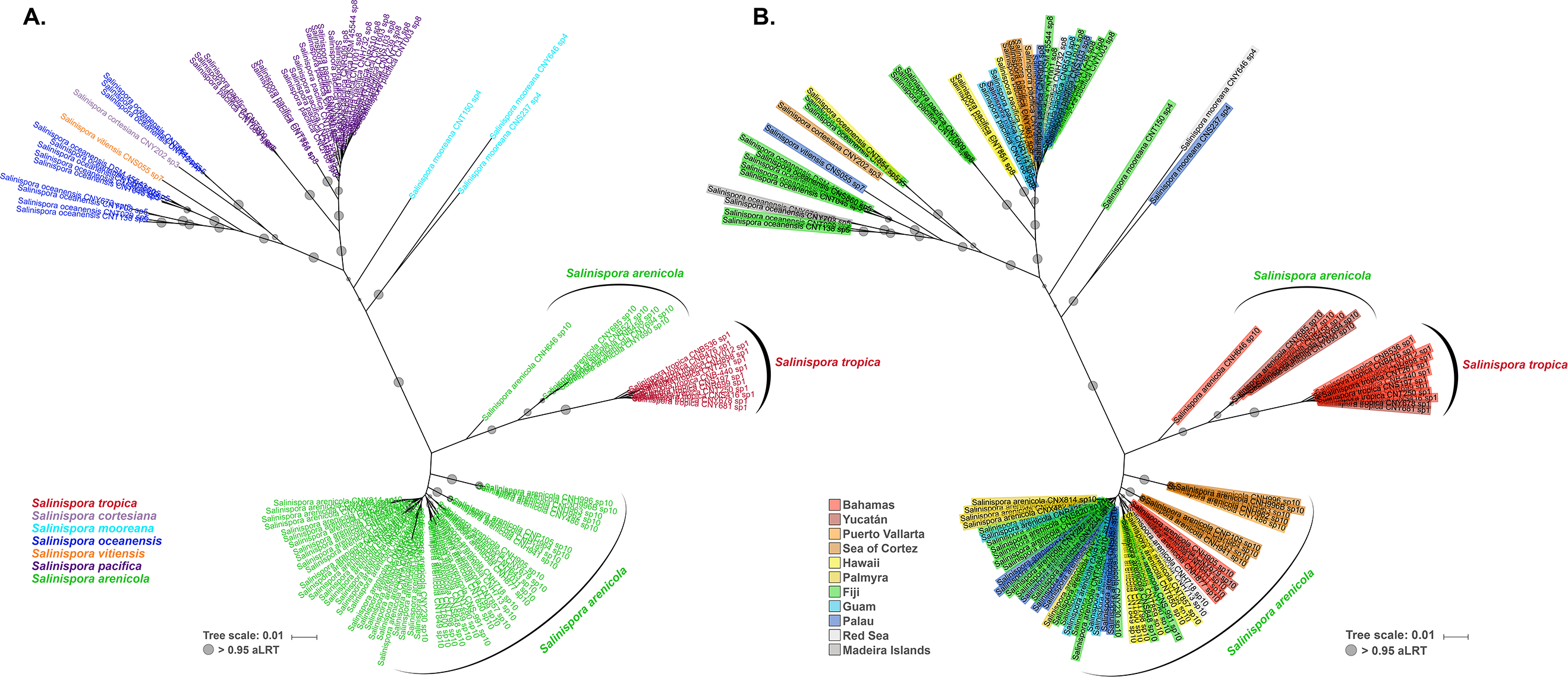
Concatenated Spt1-9 phylogeny. A) Colored by *Salinispora* species. B) Colored by *Salinispora* strain isolation location. The maximum likelihood tree was calculated in PhyML with a Smart Model Selection HIVb+G+I+F model and midpoint-rooting; branches have proportional circles representing aLRT branch support.

## Discussion

Specialized metabolites that function as signaling molecules regulate important functional traits in bacteria. However, only a small number of bacterial signaling molecules have been identified to date. This may be because they are small in size, generally produced in low yields, and often lack activity in the bioassays commonly used to guide small molecule discovery. Gamma-butyrolactones represent an important class of signaling molecules produced by Actinobacteria (Figure 1). The salinipostins, salinilactones, Sal-GBL1, and Sal-GBL2 were recently reported from the marine actinomycete genus *Salinispora* (64, 66–68) and bear structural similarities to previously characterized actinomycete γ-butyrolactones. Linkage between the biosynthesis of these compounds and the γ-butyrolactone synthase Spt9 in the salinipostin *spt* BGC led us to more broadly explore the potential for signaling molecule production by assessing the distribution of this protein among sequenced bacterial genomes. Surprisingly, we detected Spt9 homologs across 12 diverse bacterial phyla, many of which are not known to produce γ-butyrolactones (Figure 2). Despite the unexpectedly wide distribution of Spt9 homologs, only 285 of the ~25,500 currently described microbial natural products in The Natural Products Atlas (94) contain a γ-butyrolactone moiety. Of these, only 14 have been linked to their respective BGCs in the Minimum Information about a Biosynthetic Gene cluster (MIBiG) repository (95) and only four of these, including the salinipostins in *Salinispora*, A-factor and SCB1-3 in *Streptomyces coelicolor* A3(2), and lactonamycin in *Streptomyces rishiriensis*, contain an Spt9 homolog. Thus, opportunities remain to identify the products of Spt9-containing BGCs and to establish formal links between these compounds and their biosynthetic origins. Our results suggest that the production of γ-butyrolactones and related compounds may be more common that previously recognized.

A phylogenetic tree of the top 403 Spt9 homologs, including 22 AfsA homologs from experimentally characterized BGCs, showed that the associated γ-butyrolactone, γ-butenolide, and furan signaling molecules are restricted to a clade that is distinct from the majority of uncharacterized Spt9 sequences (Figure 3). The Spt9 tree also showed major incongruencies with recognized Actinobacterial and Gammaproteobacterial classification, suggesting extensive horizontal gene transfer. The genomic environments around the Spt9 homologs were diverse, suggesting the potential production of considerable chemical diversity. It remains to be seen if all of these Spt9 homologs catalyze γ-butyrolactone synthase-like reactions, especially when they are distantly related to experimentally characterized AfsA homologs. Heterologous expression of these Spt9 homologs with various precursor substrates to test if the Spt9 homologs performs the canonical AfsA condensation reaction that assembles a fatty acid ester (β-ketoacyl-DHAP ester) intermediate is a next step towards establishing their functionality (15).

Surprisingly, we discovered a large clade of Spt9 homologs in the genus *Nocardia* that shared similar operon structure with the *Salinispora spt* BGC (starred, Figure 3). To date, no small molecules isolated from *Nocardia* spp. have been linked to these BGCs. A number of differences distinguish the *Salinispora spt* BGC from the 91 *spt*-like BGCs observed both in *Nocardia* and other genera (Figure 4). Of particular note is the absence of *spt8* (a flavin-dependent oxidoreductase) outside of the genus *Salinispora* and the different gene organization, with *spt7* occurring after *spt9*, in many of the *spt-* like BGCs. Several *spt-*like BGCs also have an additional nitroreductase gene (Figure 4) between *spt9* and *spt7*, suggesting the production of a γ-butyrolactone with a reduced nitrogen or nitro functional group. Other *spt*-like BGCs lack the AMP-ligase *spt2* and the acyl carrier protein *spt4*, suggesting the products may lack the extended aliphatic sidechain observed in the salinipostins. All of these variations suggest the production of new chemical diversity and represent potential examples of BGC diversification arising in response to different selective pressures.

Also of note are the *spt*2-*spt*3 and *spt*6-*spt*9 gene fusions observed in the genera *Nocardia*, *Gordonia*, *Tsukamurella*, *Mycobacterium*, *Dietzia,* and *Streptomyces*. Both pairs of fused genes appear functional based on the maintenance of conserved functional domains (Supplementary Figure S3). *Dietzia timorensis* is the only strain with both *spt2-spt3* and *spt6-spt9* fusions and is sister to the large clade containing the other gene fusions. Protein fusions in BGCs can arise when the clustered genes are likely co-transcribed and co-translated; these fusion products can provide evidence of functional interaction and perhaps even a selective advantage over individual proteins (90, 91, 96). The gene fusions observed in the *spt*-like BGCs appear to represent the evolution of more complex, multifunctional proteins in these strains. The *spt2-spt3* and *spt6-spt9* gene fusions are similar to recently described multi-domain enzyme fusions in the desferrioxamine (*des*) BGC, where they are hypothesized to contribute to chemical diversification (96). Two additional fusiosn involving an AfsA/Spt9 homolog were observed within *trans*-AT PKS modules associated with gladiofungin and gladiostatin biosynthesis, where AfsA functions both for unprecedented offloading and butenolide formation (97, 98). Identifying unusual gene fusions such as these represents an exciting new avenue for genome-mining driven natural product discovery (96, 99).

Conservation of the *spt* BGC in the 116 *Salinispora* genomes examined here suggests it was present in a common ancestor of the genus. Yet, the *Salinispora* Spt9 phylogeny is incongruent with the established species phylogeny (Figure S4). These incongruencies are likely due to homologous recombination and horizontal gene transfer (HGT) events, the most apparent of which is represented by the clustering of the *S. tropica* Spt9 sequences within the *S. arenicola* clade. All twelve *S. tropica* Spt1-9 sequences share the same evolutionary history, providing evidence that the HGT and recombination event affected the entire BGC (Figure 5A). Co-localization of *S. tropica* and *S. arenicola* in Bahamian and Yucatán sediments provides spatial opportunities for these exchange events to occur (Figure 5B, Figure S7). Horizontal gene transfer and homologous recombination have been identified as important avenues of BGC transfer and diversification, especially in Actinobacteria (100). It remains unknown if the *S. arenicola spt* locus that appears to have replaced the ancestral version or been acquired *de novo* in *S. tropica* provides a selective advantage or affects the compounds produced. This homologous recombination and HGT event acting on an entire BGCs adds to growing evidence of gene cluster transfer between both closely and distantly related bacteria, as seen in the granaticin, coronafacoyl phytotoxins, tunicamycin, foxicin, antimycin, streptomycin, and bicyclomycin BGCs (100). BGC exchange is well documented in the genus *Salinispora* and has been linked to gene gain, loss, duplication, and divergence in lineage-specific patterns (62, 84, 101). Overall, acquisition of the *spt* BGC by *S. tropica* highlights the importance of understanding the functional roles of its products and the effects of evolutionary exchange events such as these on population and species-level dynamics.

The *spt* BGC has been linked to the production of both the salinipostins A-K (63, 64, 68), Sal-GBL1 and Sal-GB2 (68), and the salinilactones A-H (66, 67), which share structural similarities to the A-factor family of γ-butyrolactone signaling molecules.

Additionally, *S. arenicola* and *S. pacifica* produce two other AHLs that have yet to be linked to their biosynthetic genes (102). AHLs are the most common class of autoinducer signaling molecules produced by Gram-negative bacteria, thus *Salinispora* appears to employ both γ-butyrolactone and homoserine lactone signaling molecules. While further studies are needed to understand the ecological functions of the *spt* products, there is evidence that lactone signaling molecules affect microbial community organization and function (103) and can also elicit specialized metabolite production (40, 104, 105). Thus, perhaps the small molecule *spt* products regulate the expression of other biosynthetic pathways in *Salinispora*. In support of this, *spt9* genes were detected within *Salinispora* PKS and NRPS gene clusters. This is reminiscent of the methylenomycin BGC in *Streptomyces coelicolor* A3(2), which encodes the production of methylenomycin furan (MMF) signaling molecules that induce the production of the methylenomycin antibiotic (106). Additionally, we identified an *spt*-like BGC neighboring the recently identified cyphomycin PKS BGC in a Brazilian *Streptomyces* sp. ISID311 isolated from the fungus-growing ant *Cyphomyrmex* sp. (107). None of the genes in this *spt*-like BGC have been linked to cyphomycin biosynthesis (107), which suggests their small molecule products may instead have a regulatory role. Little is known about the role of signaling compounds in regulating actinomycete specialized metabolism, thus the recently reported total synthesis of salinipostin (108) and the identification of molecules from orphan *spt*-like BGCs could support future experiments designed to explore these roles.

Our results reveal unexplored biosynthetic potential related to γ-butyrolactone signaling molecules. The key biosynthetic protein sequence, Spt9, is broadly distributed among diverse bacteria and observed in a wide range of gene environments suggesting the potential for unrealized chemical diversity. Experimentally characterized γ-butyrolactone, γ-butenolide, and furan BGCs are largely restricted to the genus *Streptomyces*, yet *spt*-like BGCs are observed among bacterial genera that are not widely recognized for the production of signaling molecules. Evidence of gene fusions and gene gain/loss in the newly described *spt*-like BGCs suggest that new chemical diversity awaits discovery within this unusual class of compounds.

## Data Availability

All sequences analyzed in this paper were retrieved from publicly accessible databases including JGI IMG/MER and NCBI; all sequence accession information is included in the Supplementary Dataset S1. PCR sequences produced as part of this work can be accessed at GenBank Accession ####, listed in Table S1. Additionally, all phylogenetic tree alignment files for each tree figure are posted on the Open Science Framework page for this project, accessible at: https://osf.io/4g3mn/?view_only=6c4bdfd0afc74407bf0e6e56678a5c62

## Acknowledgements

This research was supported by the National Institutes of Health (5R01GM085770 to P.R.J. and B.S.M.), the National Science Foundation Graduate Research Fellowship Program (Grant No. DGE-1650112 to K.E.C.), and the Japan Society for Promotion of Science (JSPS Overseas Research Fellowship to Y.K.). We thank our colleagues Dr. Leesa J Klau and Dr. Henrique R. Machado for advice on analyses and figure design.

## Supplemental Figure Legends

1. S1. <u>Organization and functional annotation of the *Salinispora* salinipostin *spt* gene cluster.</u> Pfam characterizations for the nine *spt* genes (*1-9*) are given in parentheses. The structure of salinipostin A is shown.

2. S2. <u>Expanded phylogenetic tree of the top 403 Spt9 homologs (black) and 22 experimentally characterized AfsA homologs (red).</u> The RaxML maximum likelihood tree was calculated with a WAG+I+G+F ProtTest model with 500 replicates; branches are labeled with bootstrap support. Gene neighborhoods are drawn 5’ to 3’ for one representative taxa in each monophyletic clade and aligned with the Spt9 homolog in red that was used to build the tree; genes are colored by their COG function as annotated by JGI IMG/MER. Shaded rectangles indicate Actinobacterial family or Gammaproteobacterial class (see legend). Representative chemical structures are shown (γ-butyrolactones: salinipostin A from *Salinispora tropica* CNB-440, A-factor from *Streptomyces griseus*; γ-butenolide: avenolide from *Streptomyces avermitilis*; furan: methylenomycin from *Streptomyces coelicolor* A3(2)) and bracketed numbers correspond to the AfsA homologs and associated signaling molecule products in Figure 1. Stars indicate salinipostin-like BGCs.

3. S3. <u>Alignment of fused and individual Spt sequences.</u> A) Fused Spt2-3 protein sequences aligned with individual Spt2 and Spt 3 sequences from *Salinispora tropica* CNB-440 and *Streptomyces sp. PRh5.* Conserved regions are in green on the identity graph and colored by amino acid residue. The fused Spt2-3 proteins have conserved residues in the Spt2 and Spt3 functional domains (as predicted by the NCBI Conserved Domain Database tool, E-value cutoff 0.1). B) Fused Spt6-9 protein sequence from *Dietzia timorensis* ID05-A0528 aligned with individual Spt6 and Spt9 sequences from *Mycobacterium sp.* WY10, *Tsukamurella sp.* 1534, and *Nocardia carnea* NRRL B-1997. Conserved regions are in green on the identity graph and colored by amino acid residue. The fused Spt6-9 protein in *Dietzia timorensis* ID05-A0528 includes both functional domains and conserved sequence similarity to the individual Spt6 and Spt9 proteins (as predicted by the NCBI Conserved Domain Database tool, E-value cutoff 0.01).

4. S4. <u>*Salinispora* species and Spt1-9 phylogenies.</u> A) Maximum likelihood phylogenetic tree of 11 single-copy, concatenated proteins (DnaA, GyrB1, GyrB2, PyrH, RecA, Pgi, TrpB, AtpD, SucC, RpoB, and TopA) from 118 *Salinispora* genomes as reported in Ziemert et al. 2014 (84). The PhyML tree was midpoint rooted and calculated with a Smart Model Selection AIC HIVb+G+I+F amino acid model; branches are labeled with aLRT support. Taxa lacking *spt* are in black and those with the BGC are colored by species as indicated in the legend. B) Individual maximum likelihood phylogenetic trees for Spt1-9 amino acid sequences from 116 *Salinispora* strains. The PhyML trees were midpoint-rooted and calculated with a Smart Model Selection for each salinipostin protein: Spt1 (HIVb+G+I+F), Spt2 (Flu+G+I+F), Spt3 (HIVb+G+I+F), Spt4 (JTT+G), Spt5 (HIVb+G+I+F), Spt6 (HIVb+G+F), Spt7 (HIVb+G+I+F), Spt8 (HIVw+G+I+F), Spt9 (JTT+G+I+F); branches are labeled with their aLRT support. Taxa are colored by species as indicated in the legend.

5. S5. <u>*Salinispora* Spt9 phylogeny and gene cluster neighborhoods.</u> Maximum likelihood tree for Spt9 amino acid sequences from 116 *Salinispora* strains with the *spt* BGC was calculated with a JTT+I+G ProtTest model with 500 replicates in RaxML; branches are labeled with bootstrap support. Spt9 homologs in non-*Salinispora* bacteria were used as outgroups. Brackets on the left indicate genomic island (GI) locations of the *spt* BGC. Taxa are colored by species (see key). Gene neighborhoods are drawn 5’ to 3’ and aligned with the Spt9 homolog (red). Neighboring genes are colored as per MultiGeneBlast only if they share homology to genes in the *Salinispora tropica* CNB-440 *spt* BGC contig; white indicates no homology to genes in the *S. tropica* CNB-440 *spt* BGC contig. Stars indicate BGCs in which variations from the canonical *spt* BGC were observed.

6. S6. <u>Targeted PCR of *Salinispora spt* BGC variants.</u> A) The *spt* BGC in *S. arenicola* CNS-296 was observed on two contigs. Primer sets A (6F, 6dntransR) and B (6F, 6dntrans-IGR_R), which span *spt6* and the neighboring transposase gene, successfully amplified ~1400bp and ~3500bp products (gel image, lanes 1A and 1B, respectively), from *S. arenicola* CNS-296 (expected product size in parentheses). These primers failed to yield a product from *S. arenicola* CNQ-884, which possesses a contiguous *spt* BGC (gel image, lanes 2A and 2B, respectively). The light blue coverage graph indicates coverage of 8 PCR product sequences mapped to the reference genome and the identity graph shows mean pairwise identity over all mapped sequences (green: 100% identity; yellow: between 30%-99% identity; red: below 30% identity). B) Primer pair C targeting *spt6-7* amplified an ~5kb product in *S. arenicola* CNS-296 (gel images, lane 1C). Sequencing from the 5’ and 3’ ends (900bp each) mapped back to *spt6* (poorly, 67% identity) and *spt7* (99% identity) indicating an ~3.2kb insertion between *spt6* and *spt7*. Primer pair C amplified similarly sized contiguous *spt6*-7 products (~1,600bp) in both *S. arenicola* CNQ-884 (gel images, lane 2C) and *S. pacifica* CNS-143 (gel images, lane 3C). The sequences contained identical *spt6-7* sequence length and conserved domains indicating that the hypothetical gene between *spt6* and *spt7* in *S. pacifica* CNS-143 might be a genome assembly or annotation artifact. Coverage and identity graphs of mapped sequences is the same as described above.

7. S7. <u>Co-occurrence of *S. tropica* and *S. arenicola* strains.</u> Circles indicate geographic areas within the Bahamas and the Yucatán from which the 18 *Salinispora tropica* and *S. arenicola* strains that are most closely related in the Spt1-9 phylogeny (Figure S5) were isolated. The total number of strains are indicated with pie charts showing the proportion of each species; two strains are not shown as exact collection location coordinates in the Bahamas are not known for *S. tropica* strains CNT-250 and CNT-261.

## Supplemental Table Legends

Table S1. <u>NCBI GenBank Accession numbers for the *spt6-7* PCR products as described in Figure S6.</u>

## Supplementary Datasets

Dataset S1. <u>List of all sequence datasets with relevant accession information used in this paper’s analyses</u>.

Tab 1: **Fig1_AnnoTree_hits_genes**: Number of Spt9 (pfam PF03766) genome hits identified by AnnoTree across Phyla, Class, Order, Family, Genus, and Species; the proportion of all hits and the number of genomes in each clade were used to draw and scale the pie charts of Figure 2. Also listed is the Gene ID, GTDB ID, and protein sequence of all genome hits.

Tab 2: **403_spt9homolog_IMGgeneinfo**: List of all 403 Spt9 homolog sequence information (including locus tag, gene product name, genome ID, genome name, Genbank accession, amino acid sequence length, scaffold ID, scaffold external accession, scaffold length, scaffold GC%, and Pfam). Corresponds to Figure 3 and Figure S2.

Tab 3: **22_known_afsAhomologs**: List of 22 characterized AfsA homologs belonging in the gamma-butyrolactone, furan, gamma-butenolide, and ‘other’ class of signaling molecules; corresponds to Figure 1. Information includes gene locus tag, gene symbol/name, molecule linked to gene if known, genome strain, NCBI protein accession, JGI Gene ID, JGI genome ID, and gene product name.

Tab 4: **152_spt-likeBGCs**: List of all 152 Spt9 homologs identified within Spt-like BGCs and additional homologs using for building the tree in Figure 4. Gene information includes JGI gene ID, locus tag, gene product name, JGI genome ID, genome name, start coordinate, end coordinate, strand, DNA sequence length, amino acid length, scaffold ID, scaffold external accession, scaffold GC %.

Tab 5: **11_MLST_Salinispora**: List of 119 *Salinispora* species strain DnaA, GyrB1, GyrB2, Pgi, TrpB, SucC, RecA, PyrH, TopA, AtpD, RpoD JGI gene ID accessions for Figure S4-A.

Tab 6: **Spt1_Salinispora_116**: List of all 116 Spt1 *Salinispora* gene sequence information for Figure 5 and S4-B (including JGI gene ID, locus tag, gene product name, JGI genome ID, genome name, start coordinate, end coordinate, strand, DNA sequence length, amino acid sequence length, scaffold ID, scaffold external accession name, and pfam).

Tab 7: **Spt2_Salinispora_116**: List of all 116 Spt2 *Salinispora* gene sequence information for Figure 5 and S4-B (included information is the same as Spt1).

Tab 8: **Spt3_Salinispora_116**: List of all 116 Spt3 *Salinispora* gene sequence information for Figure 5 and S4-B (including JGI gene ID, locus tag, gene product name, JGI genome ID, genome name, start coordinate, end coordinate, strand, DNA sequence length, amino acid sequence length, scaffold ID, and scaffold external accession name).

Tab 9: **Spt4_Salinispora_116**: List of all 116 Spt4 *Salinispora* gene sequence information for Figure 5 and S4-B (included information is the same as Spt1).

Tab 10: **Spt5_Salinispora_116**: List of all 116 Spt5 *Salinispora* gene sequence information for Figure 5 and S4-B (included information is the same as Spt1).

Tab 11: **Spt6_Salinispora_116**: List of all 116 Spt6 *Salinispora* gene sequence information for Figure 5 and S4-B (included information is the same as Spt1).

Tab 12: **Spt7_Salinispora_116**: List of all 116 Spt7 *Salinispora* gene sequence information for Figure 5 and S4-B (included information is the same as Spt1).

Tab 13: **Spt8_Salinispora_116**: List of all 116 Spt8 *Salinispora* gene sequence information for Figure 5 and S4-B (included information is the same as Spt1).

Tab 14: **Spt9_Salinispora_116**: List of all 116 Spt9 *Salinispora* gene sequence information for Figure 5 and S4-B (included information is the same as Spt1).

## Notes

### Competing Interest Statement

The authors have declared no competing interest.

## References

1. Papenfort K, Bassler BL. 2016. Quorum sensing signal–response systems in Gram-negative bacteria. Nat Rev Microbiol 14:576–588.

2. Novick RP, Geisinger E. 2008. Quorum Sensing in Staphylococci. Annu Rev Genet 42:541–564.

3. Khokhlov AS, Tovarova II, Borisova LN, Pliner SA, Shevchenko LN, Kornitskaia EI, Ivkina NS, Rapoport IA. 1967. A-faktor, obespechivaiushchii biosintez streptomitsina mutantnym shtammom *Actinomyces streptomycini*. Dokl Akad Nauk SSSR 177:232–235.

4. Ando N, Matsumori N, Sakuda S, Beppu T, Horinouchi S. 1997. Involvement of AfsA in A-factor Biosynthesis as a Key Enzyme. J Antibiot (Tokyo) 50:847–852.

5. Lee YJ, Kitani S, Nihira T. 2010. Null mutation analysis of an afsA-family gene, barX, that is involved in biosynthesis of the γ-butyrolactone autoregulator in *Streptomyces virginiae*. Microbiology 156:206–210.

6. Sato K, Nihira T, Sakuda S, Yanagimoto M, Yamada Y. 1989. Isolation and Structure of a New Butyrolactone Autoregulator from *Streptomyces sp.* FRI-5. J Ferment Bioeng 68:170–173.

7. Hashimoto K, Nihira T, Sakuda S, Yamada Y. 1992. IM-2, a Butyrolactone Autoregulator, Induces Production of Several Nucleoside Antibiotics in *Streptomyces* sp. FRI-5. J Ferment Bioeng 73:449–455.

8. Kitani S, Yamada Y, Nihira T. 2001. Gene Replacement Analysis of the Butyrolactone Autoregulator Receptor (FarA) Reveals that FarA Acts as a Novel Regulator in Secondary Metabolism of *Streptomyces lavendulae* FRI-5. J Bacteriol 183:4357–4363.

9. Kitani S, Iida A, Izumi T aki, Maeda A, Yamada Y, Nihira T. 2008. Identification of genes involved in the butyrolactone autoregulator cascade that modulates secondary metabolism in *Streptomyces lavendulae* FRI-5. Gene 425:9–16.

10. Waki M, Nihira T, Yamada Y. 1997. Cloning and Characterization of the Gene (*farA*) Encoding the Receptor for an Extracellular Regulatory Factor (IM-2) from *Streptomyces* sp. Strain FRI-5. J Bacteriol 179:5131–5137.

11. Gräfe U, Schade W, Eritt I, Fleck WF, Radics L. 1982. A new inducer of anthracycline biosynthesis from *Streptomyces viridochromogenes*. J Antibiot (Tokyo) 35:1722–1723.

12. Gräfe U, Reinhardt G, Schade W, Eritt I, Fleck W. F., Radics L. 1983. Interspecific Inducers of Cytodifferentiation and Anthracycline Biosynthesis from *Streptomyces bikinensis* and *S. cyaneofuscatus*. Biotechnol Lett 5:591–596.

13. Zou Z, Du D, Zhang Y, Zhang J, Niu G, Tan H. 2014. A γ-butyrolactone-sensing activator/repressor, JadR3, controls a regulatory mini-network for jadomycin biosynthesis. Mol Microbiol 94:490–505.

14. Joo H-S, Yang Y-H, Lee C-S, Kim J-H, Kim B-G. 2007. Fragmentation study on butanolides with tandem mass spectrometry and its application for the screening of ScbR-captured quorum sensing molecules in *Streptomyces coelicolor* A3(2). Rapid Commun Mass Spectrom 21:764–770.

15. Kato J-Y, Funa N, Watanabe H, Ohnishi Y, Horinouchi S. 2007. Biosynthesis of y-butyrolactone autoregulators that switch on secondary metabolism and morphological development in *Streptomyces*. Proc Natl Acad Sci 104:2378–2383.

16. Efremenkova OV. 2016. A-Factor-Like Autoregulators. Russ J Bioorganic Chem 42:457–472.

17. Ceniceros A, Dijkhuizen L, Petrusma M. 2017. Molecular characterization of a *Rhodococcus jostii* RHA1 γ-butyrolactone(-like) signalling molecule and its main biosynthesis gene *gblA*. Sci Rep 7:1–13.

18. Onoprienko VV, Anisova LN, Blinova IN, Efremenkova OV, Koz’min YP, Khokhlov AS. 1983. Bioregulators of *Streptomyces coelicolor* A3(2), p. 111–112. In VII Sovetsko-indiĭskiĭ simpozium po khimii prirodnykh soedineniĭ. Tbilisi.

19. Takano E, Nihira T, Hara Y, Jones JJ, Gershater CJL, Yamada Y, Bibb M. 2000. Purification and Structural Determination of SCB1, a Gamma-Butyrolactone That Elicits Antibiotic Production in *Streptomyces coelicolor* A3(2). J Biol Chem 275:11010–11016.

20. Hsiao NH, Söding J, Linke D, Lange C, Hertweck C, Wohlleben W, Takano E. 2007. ScbA from Streptomyces coelicolor A3(2) has homology to fatty acid synthases and is able to synthesize γ-butyrolactones. Microbiology 153:1394–1404.

21. Hsiao NH, Nakayama S, Merlo ME, de Vries M, Bunet R, Kitani S, Nihira T, Takano E. 2009. Analysis of Two Additional Signaling Molecules in *Streptomyces coelicolor* and the Development of a Butyrolactone-Specific Reporter System. Chem Biol 16:951–960.

22. Sidda JD, Poon V, Song L, Wang W, Yang K, Corre C. 2016. Overproduction and identification of butyrolactones SCB1-8 in the antibiotic production superhost: *Streptomyces* M1152. Org Biomol Chem 14:6390–6393.

23. Yamada Y, Sugamura K, Kondo K, Yanagimoto M, Okada H. 1987. The structure of inducing factors for virginiamycin production in *Streptomyces virginiae*. J Antibiot (Tokyo) 40:496–504.

24. Kondo K, Higuchi Y, Sakuda S, Nihira T, Yamada Y. 1989. New Virginiae Butanolides From *Streptomyces virginiae*. J Antibiot (Tokyo) 42:1873–1876.

25. Kawachi R, Akashi T, Kamitani Y, Sy A, Wangchaisoonthorn U, Nihira T, Yamada Y. 2000. Identification of an AfsA homologue (BarX) from *Streptomyces virginiae* as a pleiotropic regulator controlling autoregulator biosynthesis, virginiamycin biosynthesis and virginiamycin M1 resistance. Mol Microbiol 36:302–313.

26. Arakawa K, Mochizuki S, Yamada K, Noma T, Kinashi H. 2007. γ-Butyrolactone autoregulator-receptor system involved in lankacidin and lankamycin production and morphological differentiation in *Streptomyces rochei*. Microbiology 153:1817–1827.

27. Arakawa K, Tsuda N, Taniguchi A, Kinashi H. 2012. The Butenolide Signaling Molecules SRB1 and SRB2 Induce Lankacidin and Lankamycin Production in *Streptomyces rochei*. ChemBioChem 13:1447–1457.

28. Yamamoto S, He Y, Arakawa K, Kinashi H. 2008. γ-Butyrolactone-Dependent Expression of the *Streptomyces* Antibiotic Regulatory Protein Gene *srrY* Plays a Central Role in the Regulatory Cascade Leading to Lankacidin and Lankamycin Production in *Streptomyces rochei*. J Bacteriol 190:1308–1316.

29. Kitani S, Miyamoto KT, Takamatsu S, Herawati E, Iguchia H, Nishitomi K, Uchida M, Nagamitsu T, Omura S, Ikeda H, Nihira T. 2011. Avenolide, a *Streptomyces* hormone controlling antibiotic production in *Streptomyces avermitilis*. Proc Natl Acad Sci 108:16410–16415.

30. Zhu J, Sun D, Liu W, Chen Z, Li J, Wen Y. 2016. AvaR2, a pseudo γ-butyrolactone receptor homologue from *Streptomyces avermitilis*, is a pleiotropic repressor of avermectin and avenolide biosynthesis and cell growth. Mol Microbiol 102:562–578.

31. Corre C, Song L, O’Rourke S, Chater KF, Challis GL. 2008. 2-Alkyl-4-hydroxymethylfuran-3-carboxylic acids, antibiotic production inducers discovered by *Streptomyces coelicolor* genome mining. Proc Natl Acad Sci 105:17510–17515.

32. Sidda JD, Corre C. 2012. Gamma-Butyrolactone and Furan Signaling Systems in *Streptomyces* Methods in Enzymology, 1st ed. Elsevier Inc.

33. Recio E, Colinas Á, Rumbero Á, Aparicio JF, Martín JF. 2004. PI Factor, a Novel Type Quorum-sensing Inducer Elicits Pimaricin Production in *Streptomyces natalensis*. J Biol Chem 279:41586–41593.

34. Matselyukh B, Mohammadipanah F, Laatsch H, Rohr J, Efremenkova O, Khilya V. 2015. N-methylphenylalanyl-dehydrobutyrine diketopiperazine, an A-factor mimic that restores antibiotic biosynthesis and morphogenesis in *Streptomyces globisporus* 1912-B2 and *Streptomyces griseus* 1439. J Antibiot (Tokyo) 68:9–14.

35. Horinouchi S. 2002. A Microbial Hormone, A-factor, as a Master Switch for Morphological Differentiation and Secondary Metabolism in *Streptomyces griseus*. Front Biosci 2045–2057.

36. Takano E. 2006. γ-Butyrolactones: *Streptomyces* signalling molecules regulating antibiotic production and differentiation. Curr Opin Microbiol 9:287–294.

37. Nishida H, Ohnishi Y, Beppu T, Horinouchi S. 2007. Evolution of γ-butyrolactone synthases and receptors in *Streptomyces*. Environ Microbiol 9:1986–1994.

38. Willey JM, Gaskell AA. 2011. Morphogenetic Signaling Molecules of the Streptomycetes. Chem Rev 111:174–187.

39. Polkade A V., Mantri SS, Patwekar UJ, Jangid K. 2016. Quorum sensing: An under-explored phenomenon in the phylum *Actinobacteria*. Front Microbiol 7:1–13.

40. Niu G, Chater KF, Tian Y, Zhang J, Tan H. 2016. Specialised metabolites regulating antibiotic biosynthesis in *Streptomyces* spp. FEMS Microbiol Rev 40:554–573.

41. Daniel-Ivad M, Pimentel-Elardo S, Nodwell JR. 2018. Control of Specialized Metabolism by Signaling and Transcriptional Regulation: Opportunities for New Platforms for Drug Discovery? Annu Rev Microbiol 72:25–48.

42. Lee KM, Lee C-K, Choi S-U, Park H-R, Kitani S, Nihira T, Hwang Y-I. 2005. Cloning and in vivo functional analysis by disruption of a gene encoding the γ-butyrolactone autoregulator receptor from *Streptomyces natalensis*. Arch Microbiol 184:249–257.

43. Healy FG, Eaton KP, Limsirichai P, Aldrich JF, Plowman AK, King RR. 2009. Characterization of γ-butyrolactone autoregulatory signaling gene homologs in the angucyclinone polyketide WS5995B producer *Streptomyces acidiscabies*. J Bacteriol 191:4786–4797.

44. Choi S-U, Lee C-K, Hwang Y-I, Kinoshita H, Nihira T. 2004. Cloning and functional analysis by gene disruption of a gene encoding a γ-butyrolactone autoregulator receptor from *Kitasatospora setae*. J Bacteriol 186:3423–3430.

45. Ichikawa N, Oguchi A, Ikeda H, Ishikawa J, Kitani S, Watanabe Y, Nakamura S, Katano Y, Kishi E, Sasagawa M, Ankai A, Fukui S, Hashimoto Y, Kamata S, Otoguro M, Tanikawa S, Nihira T, Horinouchi S, Ohnishi Y, Hayakawa M, Kuzuyama T, Arisawa A, Nomoto F, Miura H, Takahashi Y, Fujita N. 2010. Genome Sequence of *Kitasatospora setae* NBRC 14216T: An Evolutionary Snapshot of the Family *Streptomycetaceae*. DNA Res 17:393–406.

46. Aroonsri A, Kitani S, Hashimoto J, Kosone I, Izumikawa M, Komatsu M, Fujita N, Takahashi Y, Shin-ya K, Ikeda H, Nihira T. 2012. Pleiotropic Control of Secondary Metabolism and Morphological Development by KsbC, a Butyrolactone Autoregulator Receptor Homologue in *Kitasatospora setae*. Appl Environ Microbiol 78:8015–8024.

47. Intra B, Euanorasetr J, Nihira T, Panbangred W. 2016. Characterization of a gamma-butyrolactone synthetase gene homologue (*stcA*) involved in bafilomycin production and aerial mycelium formation in *Streptomyces* sp. SBI034. Appl Microbiol Biotechnol 100:2749–2760.

48. Salehi-Najafabadi Z, Barreiro C, Rodríguez-García A, Cruz A, López GE, Martín JF. 2014. The gamma-butyrolactone receptors BulR1 and BulR2 of *Streptomyces tsukubaensis*: tacrolimus (FK506) and butyrolactone synthetases production control. Appl Microbiol Biotechnol 98:4919–4936.

49. Tan G-Y, Peng Y, Lu C, Bai L, Zhong J-J. 2015. Engineering validamycin production by tandem deletion of γ-butyrolactone receptor genes in *Streptomyces hygroscopicus* 5008. Metab Eng 28:74–81.

50. Choi S-U, Lee C-K, Hwang Y-I, Kinosita H, Nihira T. 2003. γ-Butyrolactone autoregulators and receptor proteins in non-*Streptomyces* actinomycetes producing commercially important secondary metabolites. Arch Microbiol 180:303–307.

51. Du Y-L, Shen X-L, Yu P, Bai L-Q, Li Y-Q. 2011. Gamma-butyrolactone regulatory system of *Streptomyces chattanoogensis* links nutrient utilization, metabolism, and development. Appl Environ Microbiol 77:8415–8426.

52. Millán-Aguiñaga N, Chavarria KL, Ugalde JA, Letzel A-C, Rouse GW, Jensen PR. 2017. Phylogenomic Insight into *Salinispora* (Bacteria, Actinobacteria) Species Designations. Sci Rep 7:3564.

53. Román-Ponce B, Millán-Aguiñaga N, Guillen-Matus D, Chase AB, Ginigini JGM, Soapi K, Feussner KD, Jensen PR, Trujillo ME. 2020. Six novel species of the obligate marine actinobacterium *Salinispora*, *Salinispora cortesiana* sp. nov., *Salinispora fenicalii* sp. nov., *Salinispora goodfellowii* sp. nov., *Salinispora mooreana* sp. nov., … Int J Syst Evol Microbiol 70:4668–4682.

54. Mincer TJ, Jensen PR, Kauffman CA, Fenical W. 2002. Widespread and Persistent Populations of a Major New Marine Actinomycete Taxon in Ocean Sediments. Appl Environ Microbiol 68:5005–5011.

55. Jensen PR, Gontang E, Mafnas C, Mincer TJ, Fenical W. 2005. Culturable marine actinomycete diversity from tropical Pacific Ocean sediments. Environ Microbiol 7:1039–1048.

56. Mincer TJ, Fenical W, Jensen PR. 2005. Culture-dependent and culture-independent diversity within the obligate marine actinomycete genus *Salinispora*. Appl Environ Microbiol 71:7019–7028.

57. Maldonado LA, Fenical W, Jensen PR, Kauffman CA, Mincer TJ, Ward AC, Bull AT, Goodfellow M. 2005. *Salinispora arenicola* gen. nov., sp. nov. and *Salinispora tropica* sp. nov., obligate marine actinomycetes belonging to the family *Micromonosporaceae*. Int J Syst Evol Microbiol 55:1759–1766.

58. Kim TK, Garson MJ, Fuerst JA. 2005. Marine actinomycetes related to the ‘*Salinospora*’ group from the Great Barrier Reef sponge *Pseudoceratina clavata*. Environ Microbiol 7:509–518.

59. Vidgen ME, Hooper JNA, Fuerst JA. 2012. Diversity and distribution of the bioactive actinobacterial genus *Salinispora* from sponges along the Great Barrier Reef. Antonie Van Leeuwenhoek 101:603–618.

60. Jensen PR, Moore BS, Fenical W. 2015. The marine actinomycete genus *Salinispora*: a model organism for secondary metabolite discovery. Nat Prod Rep 32:738–751.

61. Feling RH, Buchanan GO, Mincer TJ, Kauffman CA, Jensen PR, Fenical W. 2003. Salinosporamide A: A highly cytotoxic proteasome inhibitor from a novel microbial source, a marine bacterium of the new genus *Salinospora*. Angew Chemie - Int Ed 42:355–357.

62. Letzel A-C, Li J, Amos GCA, Millán-Aguiñaga N, Ginigini J, Abdelmohsen UR, Gaudêncio SP, Ziemert N, Moore BS, Jensen PR. 2017. Genomic insights into specialized metabolism in the marine actinomycete *Salinispora*. Environ Microbiol 19:3660–3673.

63. Amos GCA, Awakawa T, Tuttle RN, Letzel A-C, Kim MC, Kudo Y, Fenical W, Moore BS, Jensen PR. 2017. Comparative Transcriptomics as a Guide to Natural Product Discovery and Biosynthetic Gene Cluster Functionality. PNAS 114:E11121–E11130.

64. Schulze CJ, Navarro G, Ebert D, DeRisi J, Linington RG. 2015. Salinipostins A-K, long-chain bicyclic phosphotriesters as a potent and selective antimalarial chemotype. J Org Chem 80:1312–1320.

65. Yoo E, Schulze CJ, Stokes BH, Onguka O, Yeo T, Mok S, Gnädig NF, Zhou Y, Kurita K, Foe IT, Terrell SM, Boucher MJ, Cieplak P, Kumpornsin K, Lee MCS, Linington RG, Long JZ, Uhlemann A-C, Weerapana E, Fidock DA, Bogyo M. 2020. The Antimalarial Natural Product Salinipostin A Identifies Essential α/β Serine Hydrolases Involved in Lipid Metabolism in *P. falciparum* Parasites. Cell Chem Biol 27:143–157.

66. Schlawis C, Kern S, Kudo Y, Grunenberg J, Moore B, Schulz S. 2018. Structural Elucidation of Trace Components Combining GC/MS, GC/IR, DFT-Calculation and Synthesis - Salinilactones, Unprecedented Bicyclic Lactones from *Salinispora* Bacteria. Angew Chemie Int Ed 57:14921–14925.

67. Schlawis C, Harig T, Ehlers S, Guillen-Matus DG, Creamer KE, Jensen PR, Schulz S. 2020. Extending the Salinilactone Family. ChemBioChem 21:1629–1632.

68. Kudo Y, Awakawa T, Du Y-L, Jordan PA, Creamer KE, Jensen PR, Linington RG, Ryan KS, Moore BS. 2020. Expansion of gamma-butyrolactone signaling molecule biosynthesis to phosphotriester natural products. bioRxiv 2020.10.11.335315.

69. Marchler-Bauer A, Bo Y, Han L, He J, Lanczycki CJ, Lu S, Chitsaz F, Derbyshire MK, Geer RC, Gonzales NR, Gwadz M, Hurwitz DI, Lu F, Marchler GH, Song JS, Thanki N, Wang Z, Yamashita RA, Zhang D, Zheng C, Geer LY, Bryant SH. 2017. CDD/SPARCLE: Functional classification of proteins via subfamily domain architectures. Nucleic Acids Res 45:D200–D203.

70. Mendler K, Chen H, Parks DH, Lobb B, Hug LA, Doxey AC. 2019. AnnoTree: visualization and exploration of a functionally annotated microbial tree of life. Nucleic Acids Res 47:4442–4448.

71. Altschul SF, Madden TL, Schaffer AA, Zhang J, Zhang Z, Miller W, Lipman DJ. 1997. Gapped BLAST and PSI-BLAST: a new generation of protein database search programs. Nucleic Acids Res 25:3389–3402.

72. Kearse M, Moir R, Wilson A, Stones-Havas S, Cheung M, Sturrock S, Buxton S, Cooper A, Markowitz S, Duran C, Thierer T, Ashton B, Meintjes P, Drummond A. 2012. Geneious Basic: An integrated and extendable desktop software platform for the organization and analysis of sequence data. Bioinformatics 28:1647–1649.

73. Hadjithomas M, Chen I-MA, Chu K, Huang J, Ratner A, Palaniappan K, Andersen E, Markowitz V, Kyrpides NC, Ivanova NN. 2017. IMG-ABC: New features for bacterial secondary metabolism analysis and targeted biosynthetic gene cluster discovery in thousands of microbial genomes. Nucleic Acids Res 45:D560–D565.

74. Blin K, Wolf T, Chevrette MG, Lu X, Schwalen CJ, Kautsar SA, Suarez Duran HG, de los Santos ELC, Kim HU, Nave M, Dickschat JS, Mitchell DA, Shelest E, Breitling R, Takano E, Lee SY, Weber T, Medema MH. 2017. antiSMASH 4.0—improvements in chemistry prediction and gene cluster boundary identification. Nucleic Acids Res 45:1–6.

75. Blin K, Shaw S, Steinke K, Villebro R, Ziemert N, Lee SY, Medema MH, Weber T. 2019. antiSMASH 5.0: updates to the secondary metabolite genome mining pipeline. Nucleic Acids Res 47:W81–W87.

76. Medema MH, Takano E, Breitling R. 2013. Detecting Sequence Homology at the Gene Cluster Level with MultiGeneBlast. Mol Biol Evol 30:1218–1223.

77. Szklarczyk D, Gable AL, Lyon D, Junge A, Wyder S, Huerta-Cepas J, Simonovic M, Doncheva NT, Morris JH, Bork P, Jensen LJ, Von Mering C. 2019. STRING v11: Protein-protein association networks with increased coverage, supporting functional discovery in genome-wide experimental datasets. Nucleic Acids Res 47:D607–D613.

78. Edgar RC. 2004. MUSCLE: Multiple sequence alignment with high accuracy and high throughput. Nucleic Acids Res 32:1792–1797.

79. Maddison WP, Maddison DR. 2018. Mesquite: a modular system for evolutionary analysis. 3.40.

80. Darriba D, Taboada GL, Doallo R, Posada D. 2011. ProtTest 3: fast selection of best-fit models of protein evolution. Bioinformatics 27:1164–1165.

81. Stamatakis A. 2014. RAxML version 8: A tool for phylogenetic analysis and post-analysis of large phylogenies. Bioinformatics 30:1312–1313.

82. Guindon S, Dufayard JF, Lefort V, Anisimova M, Hordijk W, Gascuel O. 2010. New algorithms and methods to estimate maximum-likelihood phylogenies: Assessing the performance of PhyML 3.0. Syst Biol 59:307–321.

83. Lefort V, Longueville JE, Gascuel O. 2017. SMS: Smart Model Selection in PhyML. Mol Biol Evol 34:2422–2424.

84. Ziemert N, Lechner A, Wietz M, Millán-Aguiñaga N, Chavarria KL, Jensen PR. 2014. Diversity and evolution of secondary metabolism in the marine actinomycete genus *Salinispora*. Proc Natl Acad Sci U S A 111:E1130–9.

85. Rambaut A. 2016. FigTree v1.4.3. 1.4.3.

86. Letunic I, Bork P. 2019. Interactive Tree Of Life (iTOL) v4: recent updates and new developments. Nucleic Acids Res 47:W256–W259.

87. Nouioui I, Carro L, García-López M, Meier-Kolthoff JP, Woyke T, Kyrpides NC, Pukall R, Klenk H-P, Goodfellow M, Göker M. 2018. Genome-Based Taxonomic Classification of the Phylum *Actinobacteria*. Front Microbiol 9:1–119.

88. Dillon SC, Bateman A. 2004. The Hotdog fold: Wrapping up a superfamily of thioesterases and dehydratases. BMC Bioinformatics 5:109.

89. Chen I-MA, Chu K, Palaniappan K, Pillay M, Ratner A, Huang J, Huntemann M, Varghese N, White JR, Seshadri R, Smirnova T, Kirton E, Jungbluth SP, Woyke T, Eloe-Fadrosh EA, Ivanova NN, Kyrpides NC. 2018. IMG/M v.5.0: an integrated data management and comparative analysis system for microbial genomes and microbiomes. Nucleic Acids Res 47:666–677.

90. Enright AJ, Iliopoulos I, Kyrpides NC, Ouzounis CA. 1999. Protein interaction maps for complete genomes based on gene fusion events. Nature 402:86–90.

91. Marcotte EM, Pellegrini M, Ng H-L, Rice DW, Yeates TO, Eisenberg D. 1999. Detecting Protein Function and Protein-Protein Interactions from Genome Sequences. Science (80-) 285:751–753.

92. Penn K, Jenkins C, Nett M, Udwary DW, Gontang EA, McGlinchey RP, Foster B, Lapidus A, Podell S, Allen EE, Moore BS, Jensen PR. 2009. Genomic islands link secondary metabolism to functional adaptation in marine Actinobacteria. ISME J 3:1193–1203.

93. Ziemert N, Podell S, Penn K, Badger JH, Allen E, Jensen PR. 2012. The natural product domain seeker NaPDoS: A phylogeny based bioinformatic tool to classify secondary metabolite gene diversity. PLoS One 7:e34064.

94. Van Santen JA, Jacob G, Singh AL, Aniebok V, Balunas MJ, Bunsko D, Neto FC, Castaño-Espriu L, Chang C, Clark TN, Little JLC, Delgadillo DA, Dorrestein PC, Duncan KR, Egan JM, Galey MM, Haeckl FPJ, Hua A, Hughes AH, Iskakova D, Khadilkar A, Lee J-H, Lee S, Legrow N, Liu DY, Macho JM, McCaughey CS, Medema MH, Neupane RP, O’Donnell TJ, Paula JS, Sanchez LM, Shaikh AF, Soldatou S, Terlouw BR, Tran TA, Valentine M, Van Der Hooft JJJ, Vo DA, Wang M, Wilson D, Zink KE, Linington RG. 2019. The Natural Products Atlas: An Open Access Knowledge Base for Microbial Natural Products Discovery ́. ACS Cent Sci 5:1824–1833.

95. Kautsar SA, Blin K, Shaw S, Navarro-Muñoz JC, Terlouw BR, van der Hooft JJJ, van Santen JA, Tracanna V, Suarez Duran HG, Pascal Andreu V, Selem-Mojica N, Alanjary M, Robinson SL, Lund G, Epstein SC, Sisto AC, Charkoudian LK, Collemare J, Linington RG, Weber T, Medema MH. 2020. MIBiG 2.0: a repository for biosynthetic gene clusters of known function. Nucleic Acids Res 48:D454–D458.

96. Chevrette MG, Gutiérrez-García K, Selem-Mojica N, Aguilar-Martínez C, Yañez-Olvera A, Ramos-Aboites HE, Hoskisson PA, Barona-Gómez F. 2020. Evolutionary dynamics of natural product biosynthesis in bacteria. Nat Prod Rep 37:566–599.

97. Niehs SP, Kumpfmüller J, Dose B, Little RF, Ishida K, Flórez L V., Kaltenpoth M, Hertweck C. 2020. Insect-Associated Bacteria Assemble the Antifungal Butenolide Gladiofungin by Non-Canonical Polyketide Chain Termination. Angew Chemie Int Ed.

98. Nakou IT, Jenner M, Dashti Y, Romero-Canelón I, Masschelein J, Mahenthiralingam E, Challis GL. 2020. Genomics-driven discovery of a novel glutarimide antibiotic from *Burkholderia gladioli* reveals an unusual polyketide synthase chain release mechanism. Angew Chemie Int Ed 10.1002/anie.202009007.

99. de Rond T, Asay JE, Moore BS. 2020. Co-Occurrence of Enzyme Domains Guides the Discovery of an Oxazolone Synthetase. bioRxiv 2020.06.11.147165.

100. Park CJ, Smith JT, Andam CP. 2019. Horizontal Gene Transfer and Genome Evolution in the Phylum *Actinobacteria*, p. 155–174. In Horizontal Gene Transfer.

101. Bruns H, Crüsemann M, Letzel A-C, Alanjary M, Mcinerney JO, Jensen PR, Schulz S, Moore BS, Ziemert N. 2017. Function-related replacement of bacterial siderophore pathways. Nat Publ Gr 12:320–329.

102. Bose U, Ortori CA, Sarmad S, Barrett DA, Hewavitharana AK, Hodson1 MP, Fuerst JA, Shaw PN. 2017. Production of N-acyl homoserine lactones by the sponge-associated marine actinobacteria *Salinispora arenicola* and *Salinispora pacifica*. FEMS Microbiol Lett 364.

103. McBride SG, Strickland M. 2019. Quorum sensing modulates microbial efficiency by regulating bacterial investment in nutrient acquisition enzymes. Soil Biol Biochem 136:107514.

104. Patteson JB, Lescallette AR, Li B. 2019. Discovery and Biosynthesis of Azabicyclene, a Conserved Nonribosomal Peptide in *Pseudomonas aeruginosa*. Org Lett 21:4955–4959.

105. Okada BK, Seyedsayamdost MR. 2017. Antibiotic dialogues: induction of silent biosynthetic gene clusters by exogenous small molecules. FEMS Microbiol Rev 41:19–33.

106. Alberti F, Leng DJ, Wilkening I, Song L, Tosin M, Corre C. 2019. Triggering the expression of a silent gene cluster from genetically intractable bacteria results in scleric acid discovery. Chem Sci 10:453–463.

107. Chevrette MG, Carlson CM, Ortega HE, Thomas C, Ananiev GE, Barns KJ, Book AJ, Cagnazzo J, Carlos C, Flanigan W, Grubbs KJ, Horn HA, Hoffmann FM, Klassen JL, Knack JJ, Lewin GR, McDonald BR, Muller L, Melo WGP, Pinto-Tomás AA, Schmitz A, Wendt-Pienkowski E, Wildman S, Zhao M, Zhang F, Bugni TS, Andes DR, Pupo MT, Currie CR. 2019. The antimicrobial potential of *Streptomyces* from insect microbiomes. Nat Commun 10:1–11.

108. Okamura H, Fujioka T, Mori N, Taniguchi T, Monde K, Watanabe H, Takikawa H. 2019. First enantioselective synthesis of salinipostin A, a marine cyclic enol-phosphotriester isolated from *Salinispora* sp. Tetrahedron Lett 60:150917.

